# Mitochondria possess a large, non-selective ionic current that is enhanced during cardiac injury

**DOI:** 10.1101/2023.11.15.567241

**Authors:** Enrique Balderas, Sandra H.J. Lee, Thirupura S. Shankar, Xue Yin, Anthony M. Balynas, Christos P. Kyriakopoulos, Craig H. Selzman, Stavros G. Drakos, Dipayan Chaudhuri

## Abstract

Mitochondrial ion channels are essential for energy production and cell survival. To avoid depleting the electrochemical gradient used for ATP synthesis, channels so far described in the mitochondrial inner membrane open only briefly, are highly ion-selective, have restricted tissue distributions, or have small currents. Here, we identify a mitochondrial inner membrane conductance that has strikingly different behavior from previously described channels. It is expressed ubiquitously, and transports cations non-selectively, producing a large, up to nanoampere-level, current. The channel does not lead to inner membrane uncoupling during normal physiology because it only becomes active at depolarized voltages. It is inhibited by external Ca^2+^, corresponding to the intermembrane space, as well as amiloride. This large, ubiquitous, non-selective, amiloride-sensitive (LUNA) current appears most active when expression of the mitochondrial calcium uniporter is minimal, such as in the heart. In this organ, we find that LUNA current magnitude increases two- to threefold in multiple mouse models of injury, an effect also seen in cardiac mitochondria from human patients with heart failure with reduced ejection fraction. Taken together, these features lead us to speculate that LUNA current may arise from an essential protein that acts as a transporter under physiological conditions, but becomes a channel under conditions of mitochondrial stress and depolarization.

## MAIN TEXT

Ion and metabolite transport across the mitochondrial inner membrane is highly regulated to prevent dissipating the electrochemical gradient responsible for ATP synthesis. Whereas a broad range of mitochondrial transporters transport metabolites across membranes electroneutrally, beyond the complexes responsible for oxidative phosphorylation the mitochondrial inner membrane remains highly impermeable to electrogenic ion transport. In fact, only a limited suite of ion channels has been identified (1).

To identify inner mitochondrial membrane (IMM) ionic currents, substantial progress has come through the application of the voltage-clamp technique to mitoplasts, mitochondria treated to disrupt the outer membrane. In the whole-mitoplast configuration, access to the matrix allows full control over the membrane potential of the entire IMM, as well as its pH and chemical composition, allowing precise measurement of ion fluxes. Such electrophysiological analysis has established well-defined ionic conductances in the mitochondrial membrane, such as UCP1 and AAC1 for protons (2, 3), the inner membrane anion channel for Cl^-^ (4), and the mitochondrial calcium uniporter for Ca^2+^ (5–7).

Ion fluxes use and deplete the electrical gradient, ΔΨ, responsible for ATP synthesis, a phenomenon known as uncoupling. The magnitude of any mitochondrial inner membrane ionic current in any given tissue reflects its unavoidable effects on ΔΨ. Thus, UCP1 proton currents have large magnitude only in brown and beige adipose tissue, where uncoupling is necessary to create the futile cycle for non-shivering thermogenesis (3, 8). Similarly, mitochondrial Ca^2+^ uniporter currents are kept at low magnitude in cells with rhythmic cytoplasmic Ca^2+^ cycling, such as mammalian cardiomyocytes or *Drosophila* flight muscle, to prevent excessive dissipation of ΔΨ (9–11). Moreover, both these portals have small single-channel conductance. Channels that have high conductance, such as the K^+^ channels, remain mostly closed, opening only under well-defined circumstances, and often only a few at a time (12, 13).

While examining the effects of deletion of the mitochondrial calcium uniporter in the heart, we happened upon a highly unusual ionic conductance. This conductance produced large currents, with mitochondrial current magnitudes greater than any described so far. It was non-selective to cations, including large organic cations such as N-methyl-D-glucamine (∼7.3 Å) (14), consistent with a large pore, yet it showed extremely strong rectification, with absent ion conduction at physiological ΔΨ. Thus, this conductance is closed during normal physiology, but its large size implies that even one or a few open channels may lead to rapid mitochondrial uncoupling. Its pharmacology did not correspond to any known mitochondrial channel species, though it was mildly inhibited by amiloride. Its activity was prevented by external Ca^2+^, corresponding to the mitochondrial intermembrane space, and its magnitude was greatest in tissues where Ca^2+^ uniporter levels were low or absent, though it was present in every tissue examined. Most intriguingly, we found that the magnitude of the current in heart was increased in multiple forms of murine and human cardiomyopathy. Taken together, our results define a *L*arge, *U*biquitous, *N*on-selective, *A*miloride-sensitive (LUNA) current.

## METHODS

### Ion channels blockers and agonists

Drugs were purchased from Sigma, Abcam, and Alomone. Dimethyl Sulfoxide (DMSO) was always kept at ≤1%. Agonists and blockers were acutely administered in the recording chamber.

### Cell culture

*NDUFB10^KO^* and isogenic control HEK-293T cells were a gift of Michael Ryan (15). Cells were grown in Dulbecco’s Modified Eagle Medium (DMEM) (ThermoFisher) with 10% fetal bovine serum.

### Animal handling and genotyping

All animal procedures were reviewed and approved by the Institutional Animal Care and Use Committee at the University of Utah. *Tfam^fl^/^fl^*mice were a gift of Nils-Göran Larsson. *Mcu^-/-^* mice were a gift of Toren Finkel and were kept in a mixed C57BL/6N and CD-1 background. *Myh6-Cre* transgenic mice were obtained from the Jackson Laboratory (Bar Harbor, ME, strain Tg [Myh6-cre]2182Mds/J, stock # 011038). Cardiac-specific knockout mice were obtained from crossing *Tfam^fl^*/*^fl^*; ^+^/^+^ X *Tfam^fl^*/*^+^*; *Myh6-Cre/*+ parents, kept on a C57BL/6J background. Phenotypes of the mice used in this manuscript have been described previously. Animals were housed under standard conditions and feed *at libitum*.

### Combined pressure-overload and myocardial infarction mouse model

Transverse aortic constriction (TAC) and coronary ligation surgery was performed as previously described (16, 17). Briefly, adult mice (4-6 month old) were treated with anesthetics and analgesics (isoflurane 2-3%). Mice were anesthetized with 2-3% isoflurane (Vet One NDC 13985-528-60), and intubated with a rodent respirator (Harvard Instruments). Analgesia was with bupivacaine (1-2 mg/kg), buprenorphine (0.05-0.1 mg/kg), and carprofen (5 mg/kg). The aorta between the brachiocephalic and left carotid was clamped using sterile TAC clips (Horizon 005200). To induce myocardial infarction (MI), a small oblique thoracotomy was performed lateral to the left intercostal line in the third costal space to expose the heart. After the opening of the pericardium, the proximal left anterior descending artery (LAD) branch of the left coronary artery was ligated with a 6-0 polypropylene suture. Sham (control) mice received the same surgical procedure without the aortic and coronary artery ligation. Animals were housed separately for recovering and were returned to group housing upon cessation of analgesic regime. After one week of surgery, we performed echocardiography on mice. Blood flow (>4000 mm/s) observed at the distal end of the TAC clip confirmed constriction and impaired left ventricular wall movement confirmed successful myocardia infarction. Echocardiography 4-weeks post-surgery showed a reduction in ejection fraction and fractional shortening, indicative of heart failure.

### Human Samples

The protocol for surgical sampling of myocardial tissue from patients was approved by the University of Utah Institutional Review Board. Written informed consent for research use of cardiac tissue was obtained prospectively from heart transplantation recipients and from next-of-kin in the case of organ donors. Four patients with advanced heart failure (New York Heart Association class III/IV) were prospectively enrolled and provided myocardial tissue samples at the time of heart transplantation. As non-failing controls, myocardial tissue was also collected from 3 heart donors without a history of cardiac disease, whose heart was not allocated for heart transplantation due to non-cardiac reasons. After excision, tissue samples were immediately snap frozen and subsequently stored at -80℃ until further processing. Patients had clinical characteristics consistent with chronic advanced heart failure with reduced ejection fraction (Table 1). Donor tissue was harvested and processed the same way as the failing hearts.

**Table 1.** Clinical characteristics for human samples.

| Condition | Description | Ejection Fraction |
| --- | --- | --- |
| HFrEF | Ischemic | 37% |
| HFrEF | Non-ischemic | 19% |
| HFrEF | Non-ischemic | 14% |
| HFrEF | Ischemic | 36% |
| Control | Unused donor | 80% |
| Control | Unused donor | 55% |
| Control | Unused donor | 62% |

### Isolation of mitochondria and preparation of mitoplasts

Crude mitochondria were isolated from HEK-293T cells. Briefly, low passage cells (P 6-7) cultured for 48 h until reaching 95% confluence in DMEM plus 10% FBS. Scraped cells were collected in ice-cold divalent-free phosphate buffered saline (DVF-PBS) centrifuged at 700x g, 4°C for 6 min. Pelleted cells were kept on ice for mitochondria isolation described in the following section.

### Mitochondria isolation from mice tissues

Mitochondria were isolated from freshly dissected mice hearts, liver, brain, and spleen. Briefly, tissues were collected from euthanized mice (80% CO_2_) and kept on ice-cold DVF-PBS. Hearts, were manually dissected to remove atria and vessels, placed on ice-cold initial medium, (in mM): 250 sucrose, 5 HEPES, 1 EGTA, and 0.1% (w/v) bovine serum albumin (BSA, pH to 7.2 with KOH; osmolality to 290-310 mOsm/L). Tissues or cells were homogenized with a Potter-Elvehjem grinder attached to an overhead stirrer (IKA, Wilmington, NC); 10-12 strokes at 250 rpm were applied. Homogenates were then centrifuged at 800x *g*, 4°C for 6 min. The supernatant was recovered and centrifuged at 8000x *g*, 4°C for 8 min. Pelleted mitochondria were re-suspended on ice-cold hypertonic mannitol (in mM): 140 sucrose, 440 D-mannitol, 5 HEPES, 1 EGTA (pH to 7.2 with KOH, osmolality 550 mOsm/L), and incubated for 8 min on ice. To break the outer membrane, mitochondria were passed through a French Press homogenizer (Sim-Aminco) at a pressure equal to 2000 p.s.i. The eluate was centrifuged at 8000 x *g*, 4°C for 8 min. Pelleted mitoplasts were re-suspended in hypertonic KCl solution (in mM): 750 KCl, 100 HEPES, 1 EGTA (pH to 7.2 with KOH), until use for whole-mitoplast patch clamping.

### Whole mitoplast voltage-clamp

The procedure was performed as described previously (9, 11). Freshly prepared mitoplasts were placed on a recording chamber filled with divalent free (DFV) KCl solution (in mM): 150 KCl, 10 HEPES, 1.5 EGTA, 1.5 EDTA. Voltage-clamp protocols were applied thought an Axopatch 200B amplifier (Molecular Devices, San Jose, CA) connected to a computer through an Analog Digital interface (Digidata 1200). After mitoplast break-in, whole-mitoplast currents were acquired at 5 kHz and filtered at 1 kHz prior to analysis with pClamp 10.3 suite. The pipette solution was composed of (in mM): K-Gluconate 150, HEPES 10, EDTA 1.5, EGTA 1.5, pH 7.2, 320-340 mOsm/L. The ionic composition was kept constant in all the experiments except for the selectivity experiments described in the main text, where K^+^ was replaced with 150 mM Na^+^, Ca^2+^, Mg^2+^, or NMDG. Unless otherwise stated all experiments were initiated recording the inner mitochondrial anionic current though IMAC, a ubiquitous mitochondrial channel. IMAC currents were always observed in DVF-KCl, and confirmed successful establishment of whole-mitoplast configuration. Deactivation of IMAC occurred when Cl^-^ ions were replaced by gluconate in the bath solution. Under DVF K-gluconate (in mM: K-Gluconate 150, HEPES 10, 1.5 EDTA and 1.5 EGTA, pH 7.2), LUNA current appeared mostly at depolarized membrane potentials (≥+80 mV). Analyses were performed in Microsoft Excel and OriginPro. For figure display purposes only, capacitive transients caused by changing levels of solutions in the bath were removed and the Simplify filter from Adobe Illustrator was applied to reduce the number of points and total figure size. Neither of these maneuvers altered the shape of the traces.

## RESULTS

### Identification of a novel mitochondrial non-selective current

While recording inner-membrane currents from mitoplasts derived from whole-animal *mitochondrial calcium uniporter* (MCU) knockout mice (*Mcu^-/-^*), we noted the presence of a novel, outwardly-rectifying cationic current (Fig. 1). In a typical protocol, we apply voltage ramps to isolated mitoplasts with an internal solution containing divalent-free (DVF, 1.5mM EDTA, 1.5mM EGTA) K^+^ gluconate and an external solution containing DVF KCl. An initial outwardly-rectifying current is carried by influx of Cl^-^ through the ubiquitous inner-membrane anion channel (IMAC, Fig. 1A). Gluconate is too large an anion to be carried by IMAC, so subsequent replacement of external DVF KCl with DVF K^+^ gluconate led to blockade of IMAC and loss of the outwards current. We noted that under these conditions, there was a slow activation over 0.5-1 minute of a large outwards current carried by K^+^ efflux. Although the current was largest at positive membrane potentials, there was measurable current at 0 mV (Fig. 1B, arrow). We subsequently named this current LUNA (*l*arge, *u*biquitous, *n*on-selective, *a*miloride-sensitive) and use the current density at +100 mV to quantify its size.

**FIGURE 1.**
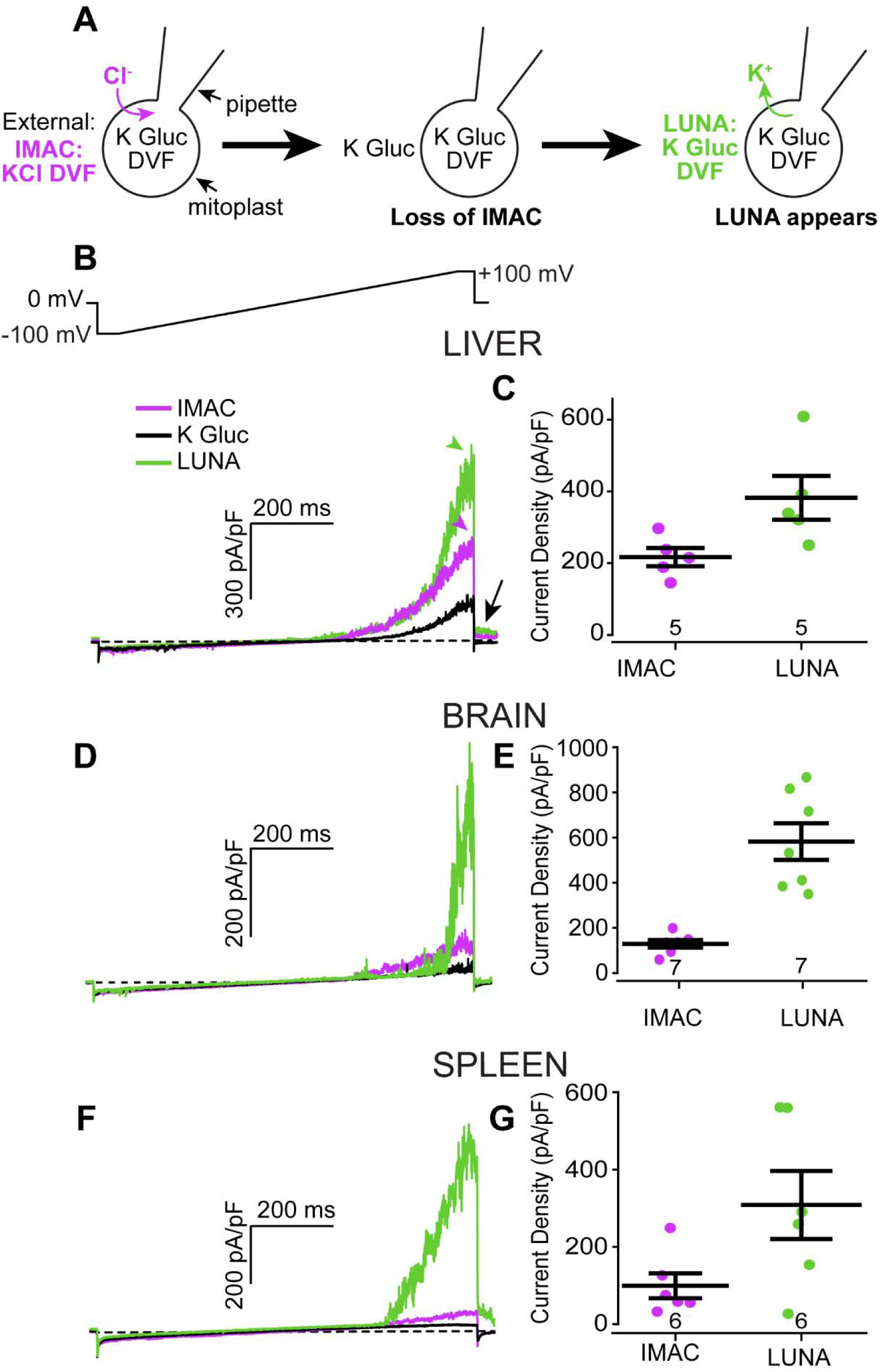
Mitochondria possess a large, divalent-sensitive outwards current. **A.** Protocol used to isolate LUNA current during whole-mitoplast electrophysiology. DVF, divalent free; IMAC, inner membrane anion channel; K Gluc, potassium gluconate. **B.** Top, voltage ramp protocol used to elicit IMAC and LUNA currents. Bottom, exemplar currents due to IMAC (purple), during washout of Cl^-^ (black), and subsequent appearance of LUNA (green) in mitoplasts isolated from *Mcu^-/-^* livers. Arrow, LUNA current is active at 0 mV holding potential. **C.** Summary of current density magnitude at +100 mV for IMAC and LUNA. N values correspond to individual mitoplasts and are reported at the bottom of the summary graph here and throughout. **D-G.** Similar to B-C, but for mitoplasts isolated from *Mcu^-/-^* whole brain and spleen.

### LUNA is ubiquitously expressed and regulated by the presence of MCU

To quantify LUNA size across different organs, we faced a complication. Under divalent-free conditions, MCU is known to permeate monovalent cations (6). Therefore, we examined tissues in *Mcu^-/-^* animals to entirely avoid potential MCU contamination. Under these conditions, LUNA was present, and of larger magnitude than IMAC in all tissues tested, comprising liver, brain, spleen, and heart (Fig. 1, 2). In contrast to IMAC, which had roughly similar magnitude across different tissues (except for liver, where IMAC was largest), there was substantial organ-specific variation in LUNA magnitude. LUNA current density was greatest in the heart (687 ± 72 pA/pF, mean ± SEM at +100 mV), followed in order by brain, liver, and spleen (545 ± 93, 382 ± 61, 308 ± 88 pA/pF, respectively). Thus, the LUNA-to-IMAC current density ratio varied from approximately two- to five-fold across these organs. This suggests that LUNA is a current carried by a separate molecular entity than IMAC.

**FIGURE 2.**
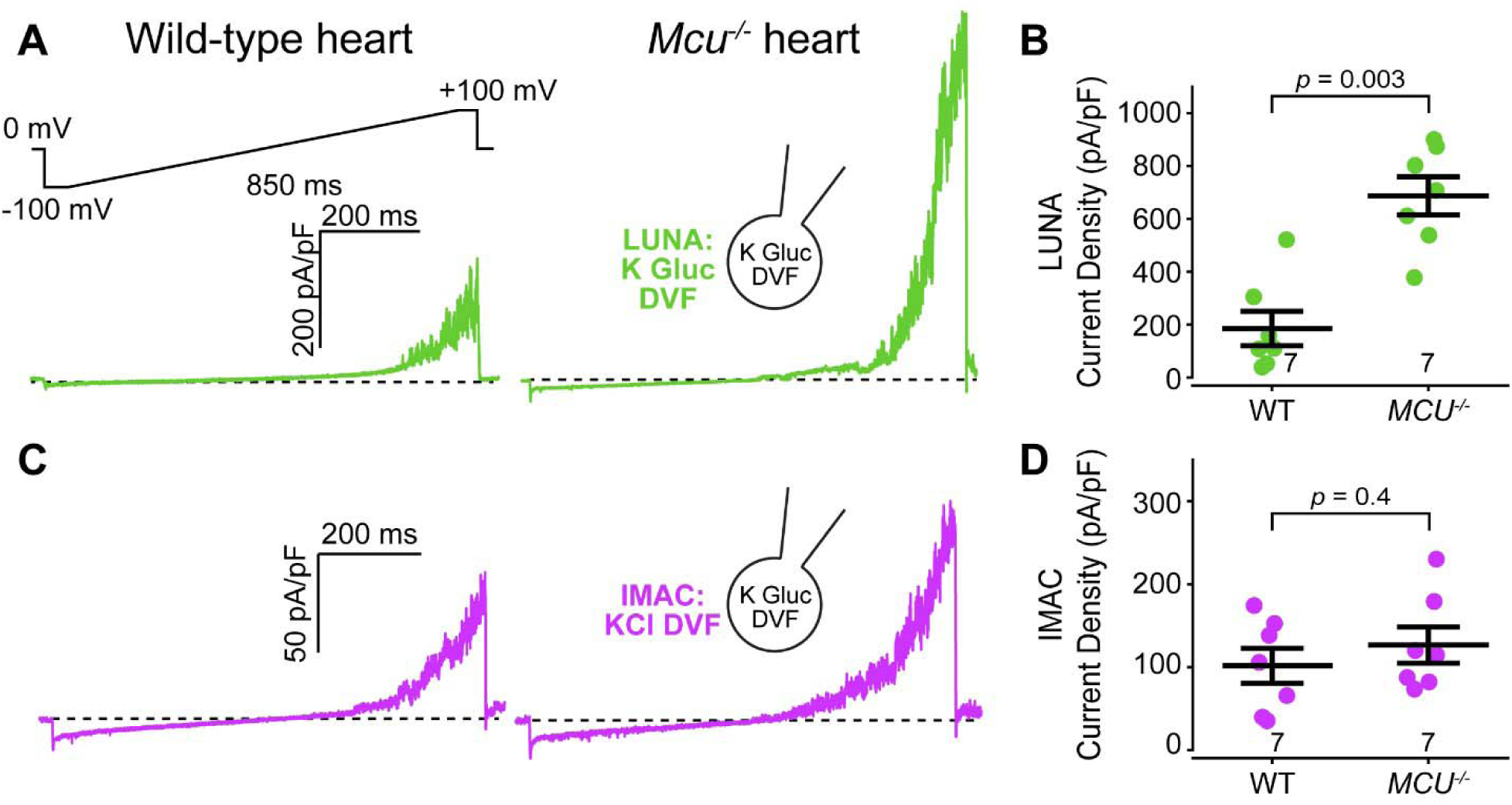
LUNA current is increased in *Mcu^-/-^* hearts. **A.** Top, voltage ramp protocol. Bottom, exemplar LUNA currents in wild-type and *Mcu^-/-^* heart mitoplasts. **B.** Summary of current density magnitude at +100 mV shows a substantial increase in the size of LUNA in *Mcu^-/-^.* **C.** Exemplar IMAC currents. **D.** Summary showing no significant change in IMAC currents in *Mcu^-/-^*. B, D. P values are for two-sided Mann-Whitney U tests here and throughout.

We were surprised that this current had not been described before. We hypothesized that knockout of MCU may have been a key intervention in making LUNA currents noticeably large. In fact, in COS-7 cell mitochondria, which have extremely large MCU currents, very limited outwards current remained after inhibiting MCU with ruthenium red (6). Conversely, in the heart, which has very low endogenous levels of MCU (18, 19), we saw the largest LUNA currents. This suggested that the magnitude of LUNA may be modulated by the presence of MCU. To examine this in more detail, we focused on the heart. To isolate LUNA in wild-type mice, we blocked monovalent currents through endogenous MCU with 1 µM ruthenium red, which does not substantially block LUNA (17% ± 5%, see Fig. 4A), and compared LUNA current density in wild-type versus *Mcu^-/-^* cardiac mitochondria (Fig. 2A-B). Surprisingly, we found LUNA current density in wild-type heart mitoplasts was 27% of its magnitude in *Mcu^-/-^*. Such regulation was specific to LUNA, as current densities for IMAC were unchanged between wild-type and *Mcu^-/-^* cardiac mitochondria (Fig. 2C-D). Taken together, this data suggests that the magnitude of LUNA is negatively regulated by MCU.

**FIGURE 3.**
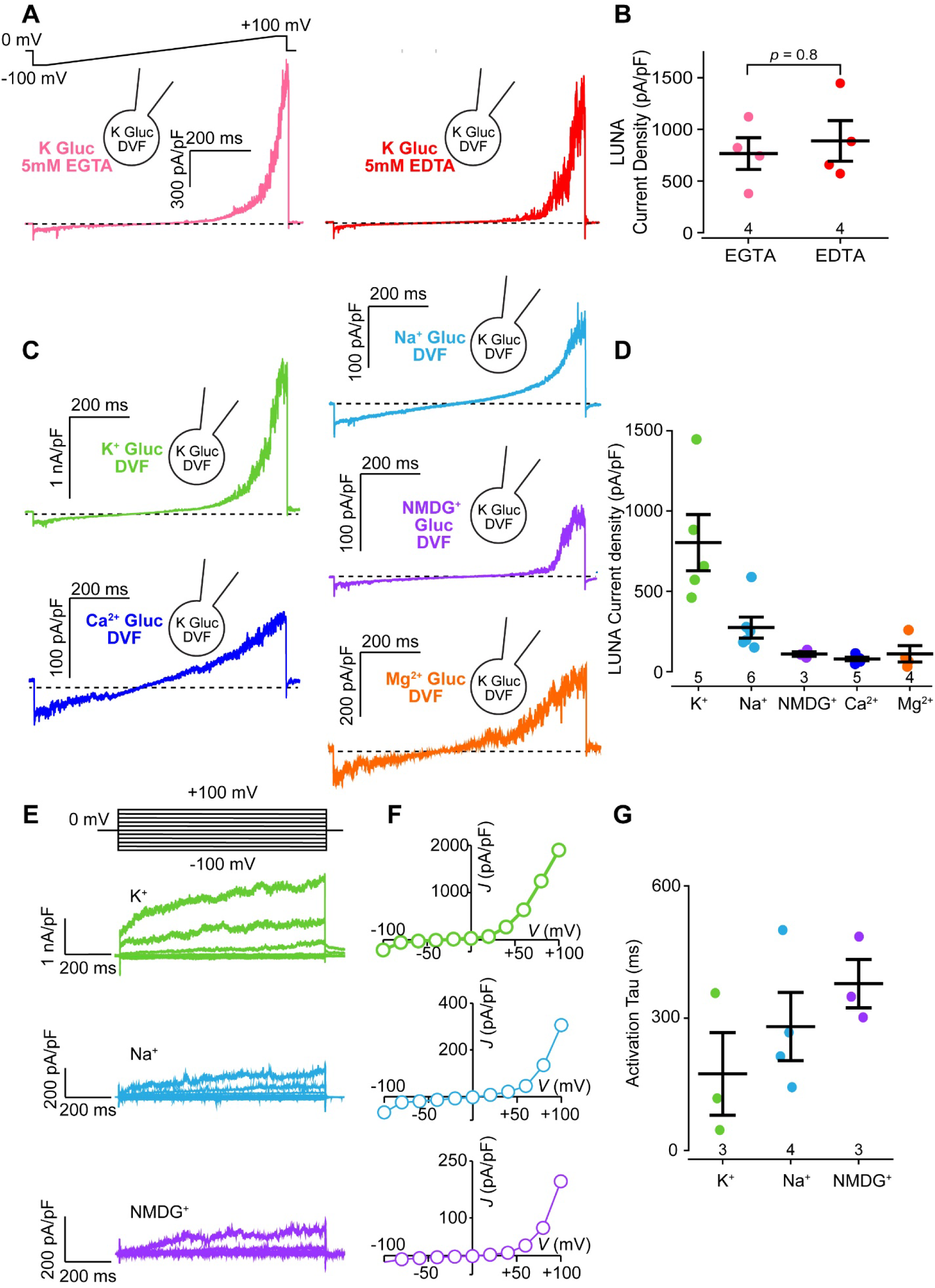
LUNA current is non-selective for cations and has slow activation kinetics. **A.** Top, voltage ramp protocol. Bottom, exemplar LUNA currents using external 5 mM EGTA or EDTA. **B.** Summary of current density magnitude at +100 mV shows no difference. **C.** Exemplar LUNA currents using different internal cations. **D.** Summary of current density magnitude at +100 mV. **E.** Step protocol to measure activation kinetics. Top, voltage step protocol. Bottom, Exemplar currents during 1 sec pulses showing slow activation. **F.** Peak current density versus step voltage traces for the corresponding exemplars in E. **G.** Summary of calculated activation time constant (tau) for each permeating cation. Data in A-G collected from *Mcu^-/-^* heart mitoplasts.

**FIGURE 4.**
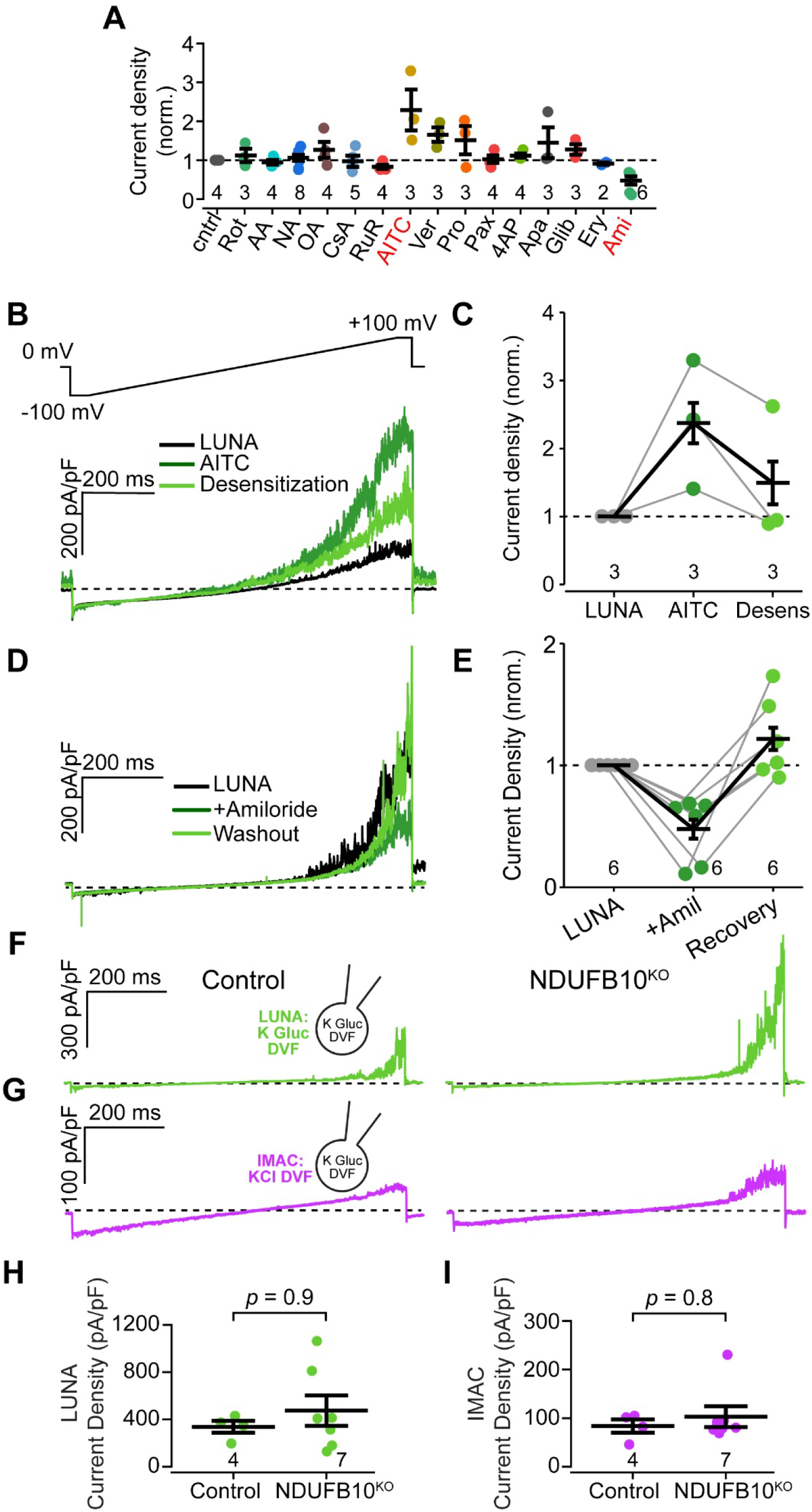
LUNA current is inhibited by amiloride, enhanced by AITC, and independent from Complex I. **A.** Summary of LUNA current density magnitude at +100 mV after application of the listed compounds. Rot, 1 µM rotenone; AA, 10 µM antimycin A; NA, 50 µM sodium azide; OA, 10 µM oligomycin A; CsA, 10 µM cyclosporine A; RuR, 1 µM ruthenium red; AITC, 10 µM allyl isothiocyanate; Ver, 10 µM verapamil; Pro, 10 µM propranolol; Pax, 1 µM paxilline; 4AP, 100 µM 4-aminopyridine; Apa, 10 µM apamin; Glib, 10 µM glibenclamide; Ery, 10 µM erythrosin B; Ami, 10 µM amiloride. **B.** Top, voltage ramp protocol. Bottom, exemplar LUNA currents at baseline (black), after addition of 10 µM AITC (dark green), and after desensitization (light green). **C.** Summary of LUNA current density magnitude at +100 mV. **D.** Exemplar LUNA currents at baseline (black), after addition of 10 µM amiloride (dark green), and after washout (light green). **E.** Summary of LUNA current density magnitude at +100 mV. **F.** Exemplar LUNA currents in control and *NDUFB10^KO^*HEK-293T mitoplasts. **G.** Exemplar IMAC currents. **H.** Summary of LUNA current density magnitude at +100 mV. **I.** Summary of IMAC current density magnitude at +100 mV. Currents in F and H were obtained in the presence of 1 µM ruthenium red to block MCU currents.

### Divalent sensitivity, selectivity, and pharmacology of LUNA

The need to use divalent-free media to maximize LUNA currents led us to examine LUNA sensitivity to external divalents in *Mcu^-/-^* heart mitoplasts. For these recordings, we used external media containing either 5mM EGTA (Ca^2+^ > Mg^2+^ selective buffer) or 5mM EDTA (Ca^2+^ and Mg^2+^ buffer) (Fig. 3A-B). LUNA was evident and of large magnitude under both conditions (EDTA: 889 ± 197, EGTA: 765 ± 153 pA/pF). Though EDTA currents were marginally higher than EGTA, there was no statistical difference between current densities in either condition.

Next, we examined LUNA selectivity. Because LUNA is strongly outwardly-rectifying, with the miniscule inwards current apparently mostly due to leak, we were unable to measure traditional permeability ratios using the Goldman-Hodgkin-Katz equation. Rather, we examine the magnitude of outwards conductance using different cations in our internal solution (K^+^, Na^+^, NMDG^+^, Ca^2+^ and Mg^2+^, all gluconate salts), while maintaining constant K^+^-gluconate in the bath solution. The relative cation conductance was K^+^ > Na^+^ > Ca^2+^ ≈ Mg^2+^ ≈ NMDG^+^ (Fig. 3C-D). Application of constant 1-sec voltage-steps revealed the kinetics of the LUNA current. There was an immediate outwards current and slower component that activated with a time constant (*τ*) that varied with the permeant ion (*τ_K_*^+^ < *τ_Na_*^+^ < *τ_NMDG_*^+^) (Fig. 3E-G). Overall, LUNA favors transport of monovalent cations with largest current amplitude and fastest activation for K^+^.

To initiate a characterization of the potential molecular carrier of LUNA, we tested a range of channel or transporter blockers. We used drugs targeting proton transporters in OXPHOS (rotenone, antimycin A, sodium azide, oligomycin A), the permeability transition pore (cyclosporine A), MCU (ruthenium red), mitochondrial anion channels (erythrosin B, propranolol), K^+^ (paxilline, 4-aminopyridine, apamin, glibenclamide), Na^+^ (amiloride), Ca^2+^ (verapamil), and TRP (allyl isothiocyanate) channels (Fig. 4A). Though most of these blockers had no significant effect on LUNA currents, we identified one potential agonist and one inhibitor of LUNA. The agonist, allyl isothiocyanate (AITC), is a broad-spectrum covalent modifier of cysteine residues, recognized for its activation of TRPA1 channels, as well as multiple non-channel targets (20). On average, 10 µM AITC enhanced LUNA current 2.3±0.4-fold upon addition to the bath medium. Notably, after reaching the maximum amplitude, LUNA currents subsequently diminished to close to their initial values, a desensitization phenomenon also seen in TRPA1 channels (Fig. 4B). The second compound identified was amiloride. When added to the bath solution, 10 µM amiloride inhibited LUNA currents by 54%±10%, with reversibility upon its removal of the solution (Fig. 4C). Though amiloride is best known as an inhibitor of Na^+^- conducting acid-sensing ion channels, it also binds to proton-pumping subunits within Complex I, as these share homology to Na^+^-H^+^ antiporters affected by amiloride (21, 22). This raised the intriguing possibility that LUNA represented a different mode of Complex I activity, behaving as a channel instead of a transporter. To test this directly, we obtained a HEK-293T cell line with deletion of a Complex I transmembrane accessory subunit, NDUFB10. In this *NDUFB10^KO^* line, there is an almost complete absence of assembled, functional Complex I, with a substantial reduction in the expression of the antiporter-like subunits as well as putative amiloride-binding subunits (15). In this cell line, despite the marked impairment in Complex I, LUNA was still present (474±119 pA/pF, Fig. 4D). Therefore, LUNA does not appear to reflect cation flux through Complex I. Nevertheless, the sensitivity of LUNA to AITC and amiloride will be useful first steps towards defining the entity conducting this novel current.

### LUNA magnitude increases in mouse models of cardiac disease

Since cardiac mitochondria had significant LUNA current and minimal MCU current, we investigated if LUNA could be regulated during disease states. Our laboratory is focused on mitochondrial cardiomyopathies, rare diseases in children due to mitochondrial mutations that lead to heart failure (11). A well-studied model for these are mice subject to cardiac-specific embryonic knockout of the nuclear-encoded transcription factor *Tfam* (*Myh6-Cre, Tfam*^lox/lox^; *Tfam* cKO). The cohort of mice we used have been described previously (9). These mice develop a cardiomyopathy, evident by postnatal day 7, driven by energetic deficiency, and leading to early demise in 3-6 weeks (11). To assay LUNA in these animals, we isolated cardiac mitoplasts from 10-14-day old *Tfam* cKO or littermate wild-type mice (3 animals each), using 1 µM ruthenium red to inhibit endogenous MCU currents. Notably, we observed that LUNA amplitude was enhanced 2.7-fold in *Tfam* cKO compared to wild-type cardiac mitoplasts (511±113, vs. 185±60 pA/pF, Fig. 5A-B). Such enhancement was specific to LUNA, as anion-selective IMAC currents were not altered (Fig. 5C-D, (11)).

**FIGURE 5.**
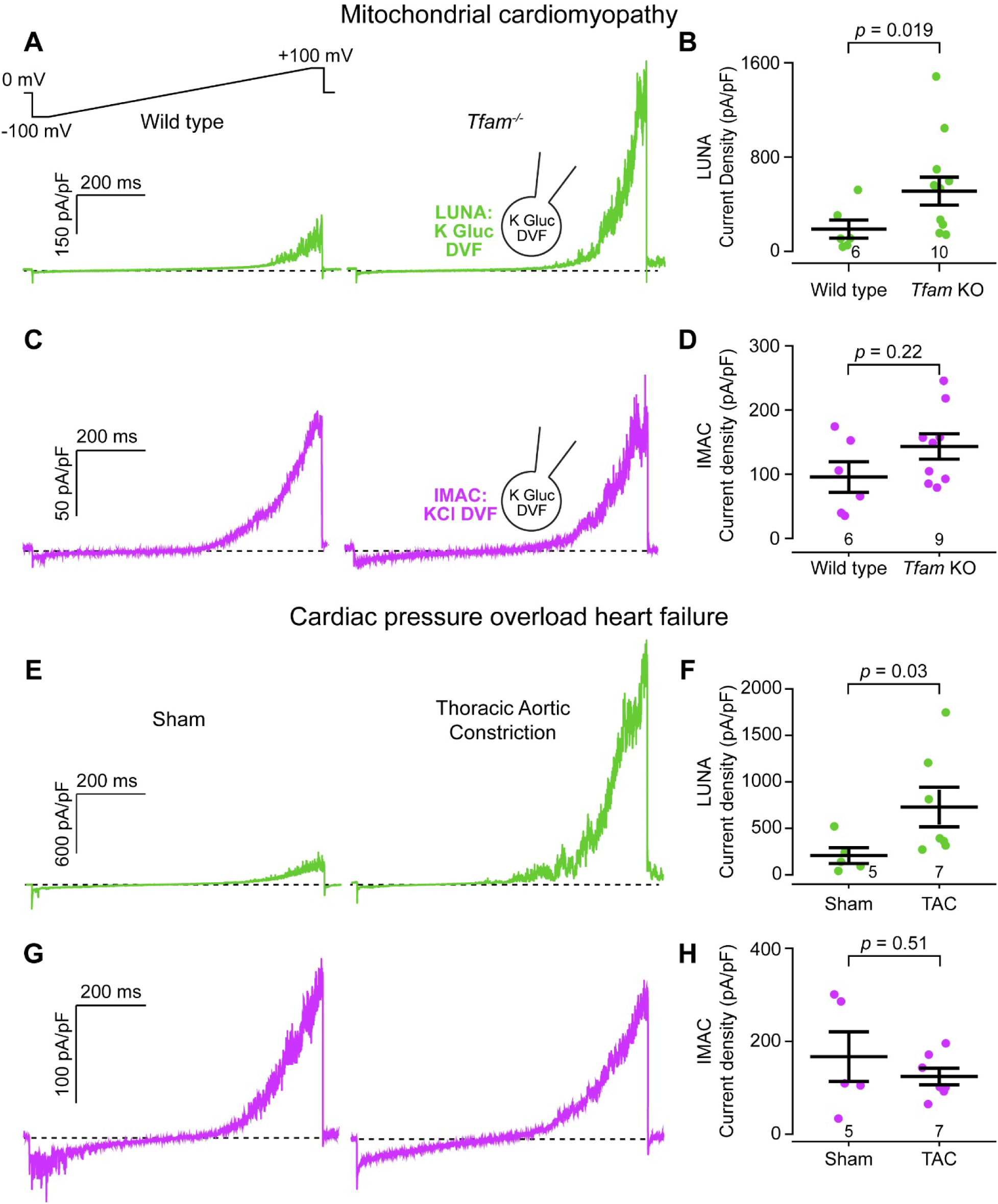
LUNA current is increased in mouse models of mitochondrial and pressure-overload cardiomyopathy. **A.** Top, voltage ramp protocol. Bottom, exemplar LUNA currents in wild-type and *Tfam*^-/-^ heart mitoplasts. **B.** Summary of LUNA current density magnitude at +100 mV, showing significant increase in *Tfam^-/-^*heart mitoplasts. **C.** Exemplar IMAC currents in wild-type and *Tfam*^-/-^ heart mitoplasts. **D.** Summary of IMAC current density magnitude at +100 mV. **E.** Exemplar LUNA currents in heart mitoplasts following sham or HF surgery procedures. **B.** Summary of LUNA current density magnitude at +100 mV, showing significant increase after HF surgery. **C.** Exemplar IMAC currents in heart mitoplasts following sham or HF surgery procedures. **D.** Summary of IMAC current density magnitude at +100 mV. Currents in A, B, E, and F were obtained in the presence of 1 µM ruthenium red to block MCU currents.

Next, to determine whether LUNA upregulation was present only during primary mitochondrial dysfunction in the *Tfam* cKO, or would occur in other models of cardiac stress, we examined a second model of cardiac disease in adult mice. We subject 4-6 month-old mice to combined pressure overload and myocardial infarction-induced heart failure (HF) or sham surgery (17). The combined insult leads to substantial heart failure 4 weeks after surgery (ejection fraction 31% ± 2%, HF surgery; 54% ± 2% sham, 3 animals each). As with *Tfam* cKO mice, we found a marked 2.6-fold increase in LUNA current density in mitoplasts derived from failing hearts (729±197 pA/pF, HF surgery; 283±137 pA/pF, Sham, Fig. 5E-F). As with *Tfam* cKO mice, the effect was specific to LUNA, with no significant change in IMAC magnitude (124 ± 16 pA/pF, HF surgery, 166 ± 47 pA/pF, Sham, Fig. 5G-H). Taken together, our data show that LUNA magnitude increases in multiple mouse models of heart failure.

### LUNA magnitude increases in human heart failure with reduced ejection fraction

Given our results in mouse models of heart disease, we next assayed LUNA in human failing hearts. We obtained cardiac mitoplasts from 4 patients with end-stage heart failure with reduced ejection fraction (HFrEF, Table 1), whose hearts had been resected during heart transplant surgery. As controls, we used cardiac mitoplasts from 3 non-failing heart donors, whose hearts could not be used for transplant due to non-cardiac reasons (e.g. size mismatch). LUNA currents were recorded with 1 µM ruthenium red to block endogenous MCU currents. As with mice, we identified robust LUNA current in non-failing donor hearts (585 ± 83 pA/pF, Fig. 6A-B). In fact, baseline cardiac LUNA was larger in humans than in mice (compare Fig. 6B to Fig. 2B). Moreover, LUNA amplitude was further increased 2.2-fold in cardiac mitoplasts from HFrEF patients, with an average amplitude that was greater than in any other condition tested in mice (heart failure or *Mcu*^-/-^) (1275 ± 182 pA/pF, Fig. 6A-B). These nanoampere-level whole-mitoplast currents are larger than any other mitochondrial conductance described in organ tissue (2, 4, 10, 11). Taken together, our results indicate that LUNA is expressed in humans and mice, and that LUNA amplitude increases during human or murine HFrEF.

**FIGURE 6.**
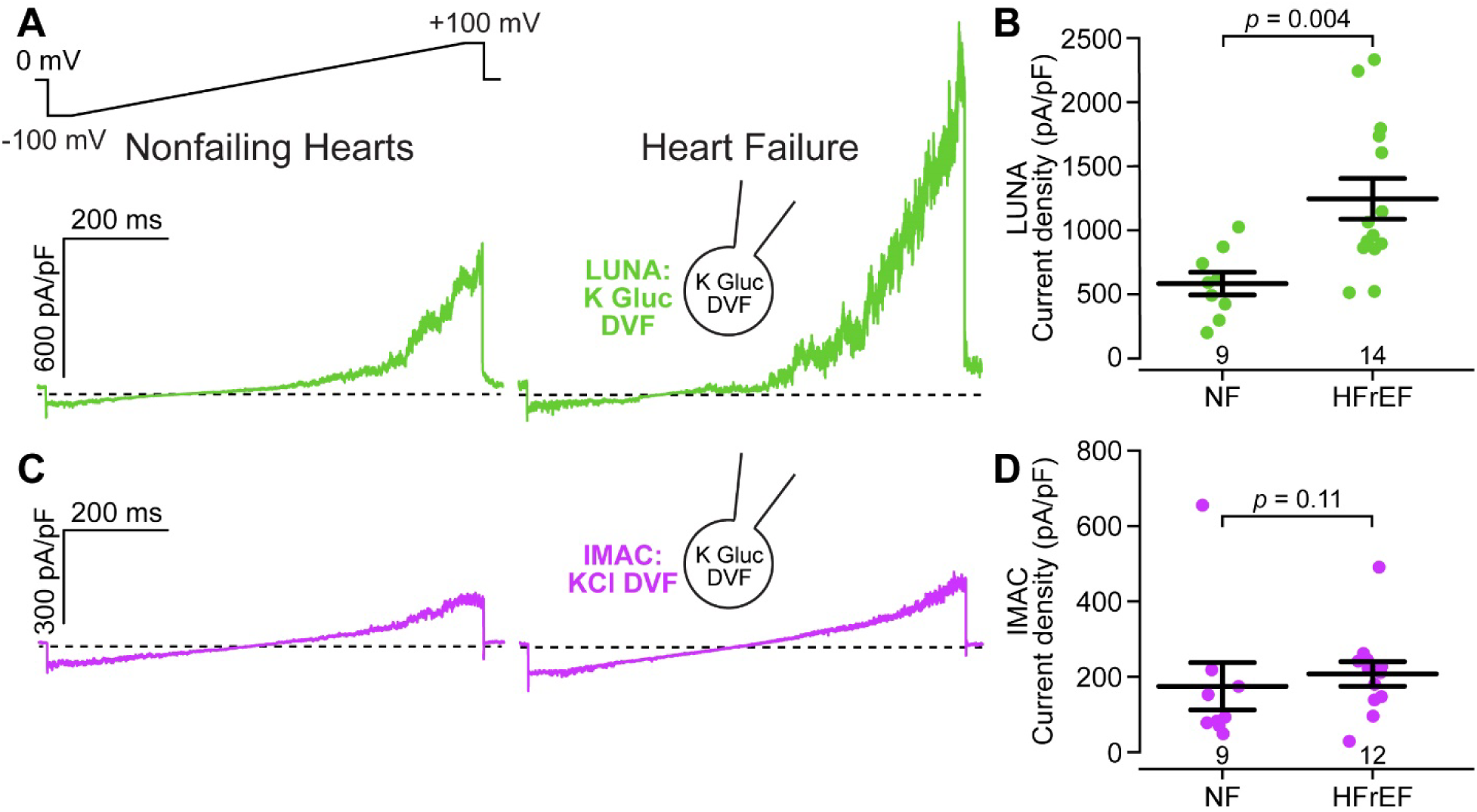
LUNA current is increased in human heart failure with reduced ejection fraction. **A.** Top, voltage ramp protocol. Bottom, exemplar LUNA currents in heart mitoplasts obtained from non-failing donors or patients with HFrEF. **B.** Summary of LUNA current density magnitude at +100 mV, showing significant increase in HFrEF. **C.** Exemplar IMAC currents in heart mitoplasts obtained from non-failing donors or patients with HFrEF. **D.** Summary of IMAC current density magnitude at +100 mV. Currents in A and B were obtained in the presence of 1 µM ruthenium red to block MCU currents.

## DISCUSSION

Here, we identify a large, non-selective, amiloride- and AITC-sensitive ionic current that appears to be expressed in every mitochondrion. LUNA current does not appear to be transported through any of the previously described mitochondrial Ca^2+^, K^+^, or Cl^-^ channels. It possesses several unusual features, described further below.

Because we have not yet defined the genetic basis for this conductance, its physiological function remains unknown. Nevertheless, its biophysical properties allow us to make several speculations. No current is evident under physiological mitochondrial transmembrane voltages (ΔΨ = -160 mV) appearing only during depolarization. This behavior is consistent with the conductance being closed during normal mitochondrial function. Because the electron transport chain expends substantial energy in producing the negative voltages needed for driving ATP synthesis through the F1-Fo-ATP synthase, channels open at these voltages would create futile cycles, of which only UCP1 is well-described for brown-fat thermogenesis. It is therefore possible that at physiological voltages, the channel conducting LUNA performs some other function. We speculate that it may be a transporter under normal conditions, as there are several examples of plasma membrane proteins with dual function (23). Members of the TMEM16 family can act as lipid-scramblases or anion-channels, with TMEM16A possessing both activities (24). Similarly, members of the CLC family can be either Cl^-^ channels or Cl^-^/H^+^ antiporters (25). The Na^+^/K^+^ ATPase can be turned into an ion channel by the reef coral poison palytoxin (26). And there are a range of transporters, such as those for glutamate, which can be induced to have channel-like conductances (27). If LUNA indeed arises from a transporter, it will be particularly notable in having transport and channel activities readily evident and controlled primarily by the direction of the membrane gradient.

Nevertheless, even if it is a transporter under physiological conditions, there is measurable current at potentials more negative than 0 mV, suggesting the channel may become active under during cellular stimulation or stress. Even opening of one or a few LUNA-carrying channels may alter mitochondrial ΔΨ and function, as seen for other large-conductance channels (12, 28). By permitting controlled ion passage across the inner mitochondrial membrane exclusively under depolarization, this channel could significantly impact the electrochemical potential gradient crucial for ATP synthesis and reactive oxygen species production. Furthermore, it appears the channel carrying LUNA is significantly inhibited by Ca^2+^. Beyond requiring Ca^2+^ chelation to fully activate the current, LUNA magnitude is greatest in mitochondria where MCU expression is limited, such as the heart, or after MCU knockout. Finally, the current is clearly regulated by stress, as its magnitude increases in multiple forms of cardiac injury in humans and mice, though it remains to be established if it contributes directly to cardiac pathophysiology. Taken together, our data suggests that membrane potential, Ca^2+^, and stress regulate the LUNA current, and this current may lead to novel adaptations to energetic demands.

Another unusual feature of the LUNA current is that, despite its strong outward rectification, when open it permeates large cations. We find that LUNA permeates NMDG, which has a mean diameter of ∼7.3 Å (14). Large-pore channels such as connexins and pannexins, which mediate the transport of metabolites between cells or their export into the extracellular space, typically conduct in both directions. Of note, to induce LUNA current, we had to chelate external divalents. This is somewhat reminiscent of the phenomenon of pore dilation in TRP channels. In certain TRP channels, prolonged exposure to agonists induces increased permeability to large cations such as NMDG. For TRPA1, TRPV1, and TRPV3, such pore dilation is enhanced substantially by chelating divalents (29, 30), suggesting that ion binding at a site close to the pore constrains its ability to dilate. However, even for these channels, pore dilation is a phenomenon producing ion conduction in both directions. The poorly-rectifying currents conducted by these large-pore channels stands in stark contrast to the highly outward-rectifying LUNA current.

This collection of unusual features and its ubiquitous expression across tissues suggest that the channel underlying LUNA may be essential for mitochondrial function. Identifying the protein encoding the channel will help clarify how LUNA fits into mitochondrial activities. Unfortunately, the gap between discovery of well-defined mitochondrial ionic conductances and identification of the corresponding channels stretched to decades for MCU and mitochondrial K^+^ and Cl^-^- channels, though the use of novel screening technologies and integrative genomics may accelerate this process. Investigating the intricate connections between this newfound ionic current, cellular energy dynamics, and physiological reactions to stress presents an exciting prospect for deepening our insights into mitochondrial physiology.

## ACKNOWLEDGEMENTS

The authors are grateful to the donor families for their generosity, and DonorConnect (https://www.donorconnect.life), Salt Lake City, Utah, for facilitating the work of our research team members acquiring myocardial tissue in the operating rooms of several hospitals of the Intermountain West. We thank Toren Finkel for gift of *Mcu^-/-^* mice, Nils-Göran Larsson for gift of floxed *Tfam* mice, Vamsi Mootha for the gift of *MCU* KO cells, and Michael Ryan for the gift of *NDUFB10* KO cells.

This work was supported by funding from grants by the National Institutes of Health HL165797, HL141353 (D.C.), HL165806 (S.L.), and the Nora Eccles Treadwell Foundation (D.C.). The content is solely the responsibility of the authors and does not necessarily represent the official views of the National Institutes of Health.

